# Dinoflagellate symbionts escape vomocytosis by host cell immune suppression

**DOI:** 10.1101/864579

**Authors:** Marie R. Jacobovitz, Sebastian Rupp, Philipp A. Voss, Sebastian G. Gornik, Annika Guse

**Author notes:** contributed equally (Marie R. Jacobovitz, Sebastian Rupp). corresponding author A.G.

## Abstract

Emergence of the symbiotic lifestyle fostered the immense diversity of all ecosystems on Earth, but symbiosis plays a particularly remarkable role in marine ecosystems. Photosynthetic dinoflagellate endosymbionts power reef ecosystems by transferring vital nutrients to their coral hosts. The mechanisms driving this symbiosis, specifically those which allow hosts to discriminate between beneficial symbionts and pathogens, are not well understood. Here, we uncover that host immune suppression is key for dinoflagellate endosymbionts to avoid elimination by the host using a comparative, model systems approach. Unexpectedly, we find that the clearance of non-symbiotic microalgae occurs by non-lytic expulsion (vomocytosis) and not intracellular digestion, the canonical mechanism used by professional immune cells to destroy foreign invaders. We provide evidence that suppression of TLR signalling by targeting the conserved MyD88 adapter protein has been co-opted for this endosymbiotic lifestyle, suggesting that this is an evolutionarily ancient mechanism exploited to facilitate symbiotic associations ranging from coral endosymbiosis to the microbiome of vertebrate guts.

## Main Text

### Summary

Symbiotic interactions appear in all domains of life and are key drivers of adaption and evolutionary diversification. A prime example is the endosymbiosis between corals and eukaryotic, photosynthetic dinoflagellates, which the hosts take up from the environment by phagocytosis. Once intracellular, endosymbionts transfer vital nutrients to the corals, a process fundamental for survival in nutrient-poor environments (Muscatine, 1990; Yellowlees et al., 2008). A central question is how endosymbionts circumvent the hosts’ defensive strategies to persist intracellularly and conversely, how hosts prevent invasion by non-symbiotic microorganisms. Here, we use a comparative approach in the anemone model *Exaiptasia pallida* (commonly *Aiptasia*) (Hambleton et al., 2019; Tolleter et al., 2013; Weis et al., 2008) to address this. Using RNA-seq, we show that symbiont phagocytosis leads to a broad scale, cell-specific transcriptional suppression of host innate immunity, but phagocytosis of non-symbiotic microalgae does not. We show that targeting MyD88 to inhibit canonical TLR signalling promotes symbiosis establishment. Using live imaging and chemical perturbations, we demonstrate that non-symbiotic microalgae are cleared from the host by ERK5-dependent vomocytosis (non-lytic expulsion) (Alvarez and Casadevall, 2006; Gilbert et al., 2017; Ma et al., 2006), and that immune suppression is key for symbiont persistence and for establishment of an intracellular LAMP1 niche. We propose that expulsion is a fundamental mechanism of ancient innate immunity used to eliminate foreign cells; a process that is subverted by symbionts in the endosymbiotic association that forms the foundation of coral reef ecosystems.

## Results & Discussion

### Phagocytosis of microalgae from the environment is primarily indiscriminate in Aiptasia larvae

The long-term pairings between cnidarian hosts and eukaryotic dinoflagellate symbionts (family Symbiodinaceae) (LaJeunesse et al., 2018) are evolutionarily ancient and highly specific (Coffroth et al., 2010; Hambleton et al., 2014; LaJeunesse et al., 2004); yet cnidarians also phagocytose inert beads (Bucher et al., 2016), associate transiently with heterologous symbiont strains (Dunn and Weis, 2009; Hambleton et al., 2014; Matthews et al., 2017; Wolfowicz et al., 2016), and with *Chromera velia*, a chromerid microalga that is closely related to apicomplexan parasites (Cumbo et al., 2013; Mohamed et al., 2018). Unlike true symbionts, however, these non-symbiotic microalgae fail to engage in lasting relationships with their host.

To gain insight into the mechanisms that support intracellular maintenance of symbionts and removal of non-symbiotic microorganisms, we exploited the ability of naturally aposymbiotic (symbiont-free) *Aiptasia* larvae to phagocytose symbionts from the environment (Bucher et al., 2016; Hambleton et al., 2014; Wolfowicz et al., 2016). We screened seven distinct microalgae from four phyla (Fig. 1a) and found that, in addition to the dinoflagellate symbiont *Breviolum minutum* (strain SSB01) (LaJeunesse et al., 2018; Xiang et al., 2013), *Chromera velia, Isochrysis sp.*, *Chlorella sp., Dunaliella salina*, *Chlamydomonas parkeae*, *Nannochloropsis oculata*, and *Microchloropsis gaditana* (Fig. 1a) are all phagocytosed by the endodermal cells of *Aiptasia* larvae (Fig. 1b, Fig. S1). For further comparative analyses we chose *C. velia* because of its interesting phylogenetic position, as well as *N. oculata* and *M. gaditana*, which have not previously been observed in association with cnidarians. We confirmed that all three microalgae are fully internalised using confocal microscopy (Fig. 1c).

**Figure 1.**
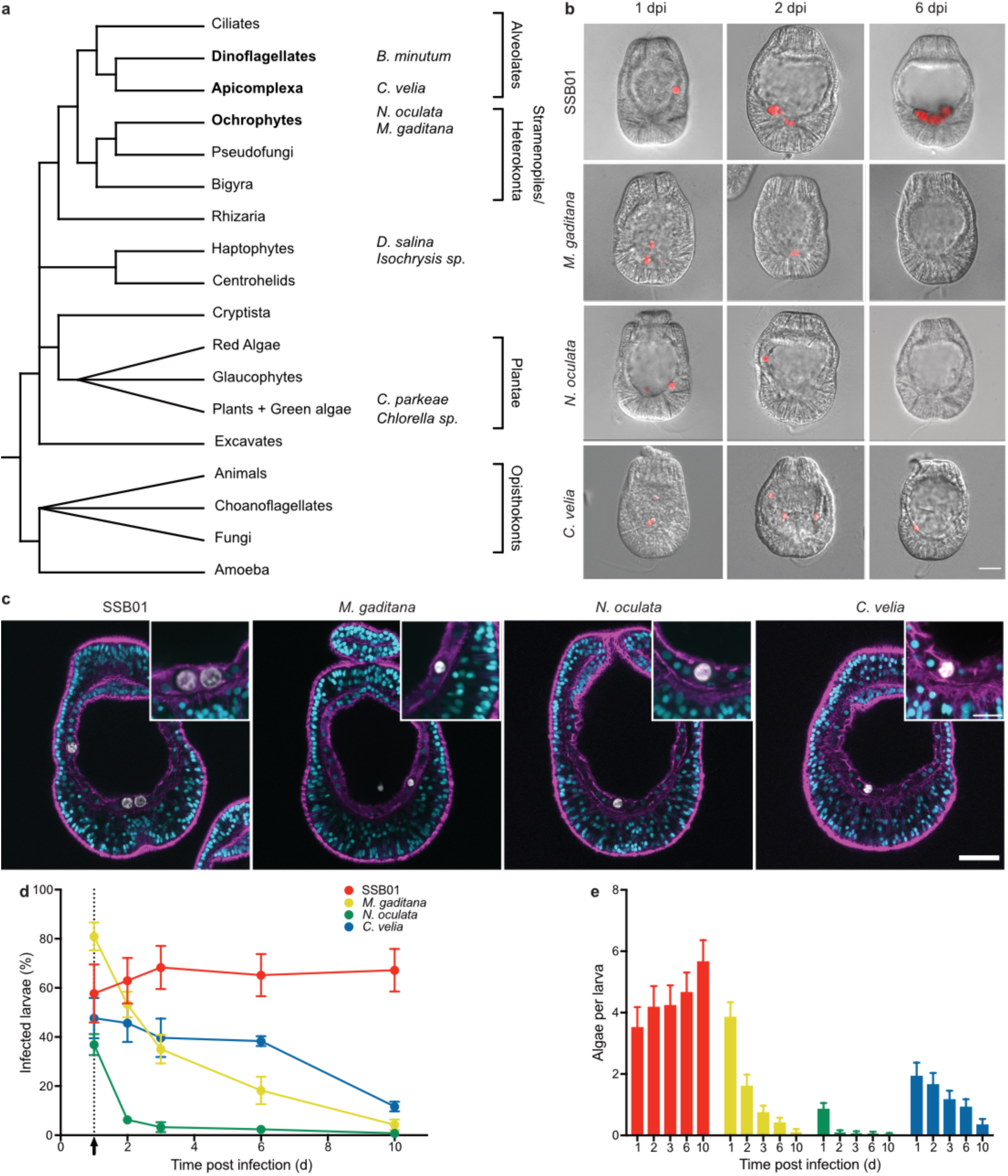
*Aiptasia* larvae as a comparative system to dissect symbiont maintenance. **a** Phylogenetic tree adapted from (Keeling and Burki, 2019). The four microalgae used for the comparative analysis belong to the SAR supergroup: *B. minutum* strain SSB01 (symbiont) belongs to the phylum Dinoflagellata, *C. velia* belongs to the phylum Apicomplexa, and both *M. gaditana* and *N. oculata* are members of Ochrophyta. The additional microalgae screened belong to two different phyla: *D. salina* and *Isochrysis sp.* - Haptophyta and *C. parkaea* and *Chlorella sp*. - Chlorophyta (images in Fig. S1). **b** *Aiptasia* larvae 1-, 2-, and 6-day(s) post infection (dpi) with SSB01, *M. gaditana*, *N. oculata*, and *C. velia*, as indicated. Larvae were infected at 4-6 days post fertilisation (dpf) for 24 hours and were washed into fresh FASW. Images are DIC and red autofluorescence of algal photosynthetic pigments, scale bar represents 25 µm. **c** All microalgae, SSB01, *M. gaditana*, *N. oculata*, and *C. velia*., are taken up in the endodermal tissue. Autofluorescence of algal photosynthetic pigment (white), Hoechst-stained nuclei (cyan), phalloidin-stained F-actin (magenta). Scale bar for whole larva represents 25 µm and scale bar for close up represents 10 µm. **d** Percentage of *Aiptasia* larvae infected after exposure to 1.0 × 10^5^ cells/ml SSB01, *M. gaditana*, *N. oculata*, *or C. velia* for 24 hours followed by a washout into fresh FASW (indicated by the arrow and dotted line). Error bars indicate mean ± SEM of 4 independent replicates. **e** Average number of algal cells/larva after exposure to 1.0 × 10^5^ cells/ml of SSB01, *M. gaditana*, *N. oculata*, *or C. velia* for 24 hours after which larvae were washed into fresh FASW. Error bars are mean ± SEM of 4 independent replicates.

To compare long-term host tolerance between symbiotic microalgae and non-symbiotic microalgae, we quantified infection rates and the number of intracellular microalgae over time. We found that the proportion of larvae containing symbionts (SSB01) remains relatively constant, even after removal of microalgae from the environment at 24 hours post infection (hpi) (Fig. 1d). In contrast, the number of larvae infected with the other algal species decreased over time. The most rapid reduction occurred with *N. oculata*, followed by *M. gaditana*, and a more gradual decrease with *C. velia*. While the number of symbiont cells increased due to proliferation within the host (Fig. 1e) (Hambleton et al., 2014), the mean algal cell count per larva for all the other microalgae decreased (Fig. 1e). Taken together, our results indicate that phagocytosis of microalgae is largely indiscriminate, suggesting that decisive symbiont-selection mechanisms occur after uptake, and that host clearance response varies between the distinct algal types.

### Symbiosis establishment broadly suppresses host cell immunity at the transcriptional level

As the sister group to bilaterians, cnidarians have a complex innate immune system that is implicated in recognising pathogens and managing beneficial microorganisms, including endosymbionts. Previous transcriptomic analyses comparing aposymbiotic to symbiotic *Aiptasia* larvae and adult animals revealed several immune-related genes to be downregulated in response to symbiosis (reviewed in Mansfield and Gilmore, 2018). Specifically, symbionts have been proposed to induce immune suppression through NF-κB downregulation, possibly via TGFβ-signalling (Berthelier et al., 2017; Detournay et al., 2012; Mansfield et al., 2019, 2017). In contrast, heterologous microalgae are thought to activate host innate immunity leading to their subsequent removal (Mansfield et al., 2019; reviewed in Mansfield and Gilmore, 2018; Mohamed et al., 2018). Selection of true endosymbionts, however, as well as rejection of non-symbiotic microalgae, occurs at the level of the individual cell. We currently lack direct evidence whether and/or how symbiont uptake specifically modulates host cell expression of innate immunity related genes, and how the transcriptional response induced by symbionts compares to that induced by non-symbiotic microalgae. To address this, we directly compared the host cell transcriptional response post-phagocytosis of either symbionts or *M. gaditana* cells, which are lost rapidly, yet remain intracellular long enough for analysis (Fig. 1d), using our recently developed cell-specific RNA-seq approach (Fig. S2a) (Voss et al., 2019). We infected *Aiptasia* larvae with *M. gaditana* for 24-48 h, dissociated infected larvae, and sequenced groups of 8-12 endodermal cells that contained *M. gaditana* cells and aposymbiotic cells from *M. gaditana*-infected larvae. We compared *M. gaditana-*containing endodermal cells and aposymbiotic endodermal cells to endodermal cells containing symbionts and aposymbiotic cells from symbiotic larvae, as well as cells from aposymbiotic larvae (Fig. S2a) (Voss et al., 2019). Using this cell-type specific approach, we found that only cells containing symbionts are clearly distinct in their overall gene expression when compared to all other cell types (Fig. 2a, Fig. S2b) (Voss et al., 2019). In contrast, *M. gaditana*-containing cells are basically indistinguishable at the transcriptional level from aposymbiotic cells (Fig. 2a, Fig. S2b), suggesting that the recognition and removal of non-symbiotic microalgae by the host cell may predominantly be regulated at the post-transcriptional level.

**Figure 2.**
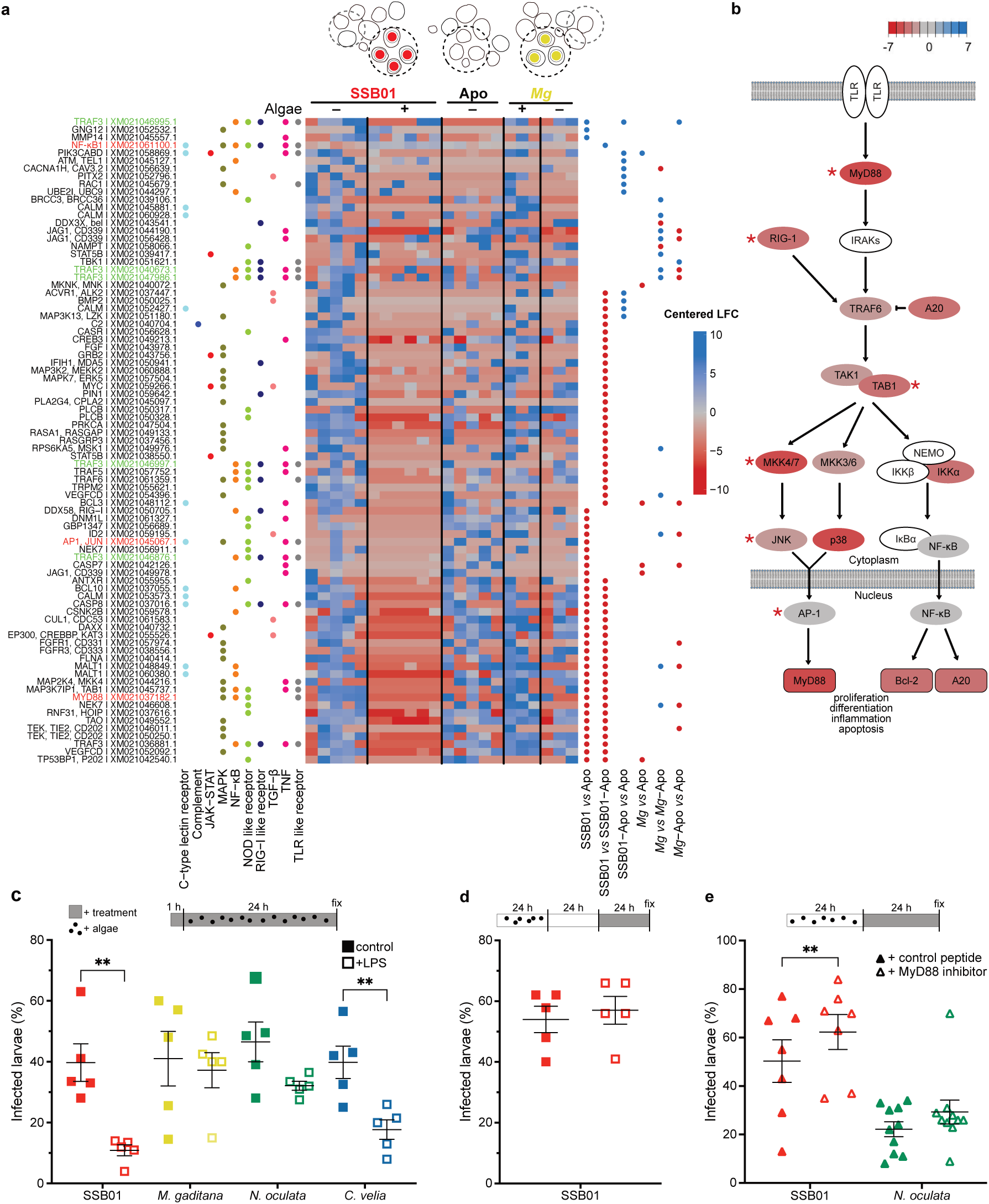
Modulation of host innate immunity is involved in symbiosis establishment. **a** Several genes in immunity related pathways are differentially downregulated in SSB01 (symbiont) containing cells, but not in *M. gaditana* (Mg) containing cells, or aposymbiotic cells from infected larvae (values indicated centred log fold change according to Deseq2 with downregulation in red and upregulation in blue). The heatmap shows all differentially regulated genes SSB01 vs Apo, SSB01 vs SSB01-Apo, SSB01-Apo vs Apo, Mg vs Apo, Mg vs Mg-Apo, Mg-Apo vs Apo within the following KEGG pathways: C-type lectin receptor signalling pathway (ko04625), Complement and coagulation cascades (ko04610), JAK-STAT (ko04630), MAPK (ko04010), NF-κB (ko04064), NOD-like receptor (ko04621), RIG-I-like receptor (ko04622), TGFβ (ko04350), TNF (ko04668), and TLR-like receptor (ko04620). Significantly differentially expressed genes compared between populations of single cells are indicated with blue (up-regulated) or red (down-regulated) dots. Gene names in green indicate multiple transcripts annotated as the same gene (TRAF3) and gene names in red are those of special interest as mentioned throughout the text. KEGG annotation was automated based on homology. **b** Simplified TLR pathway according to KEGG annotations (genes in ‘white’ could not be identified in *Aiptasia*) representing differential gene expression between symbiotic cells *vs*. aposymbiotic cells from naïve larvae. Asterisks indicate statistically significant changes. **c** Percentage of SSB01 (symbiont)-infected *Aiptasia* larvae is reduced (*p*=0.0057, t=5.391) when larvae are pre-treated for 1 hour with LPS, followed by a 24-hour exposure to microalgae and LPS. Percentage of *C. velia*-infected larvae is also reduced (*p*=0.0034, t=6.205), but to a lesser degree, but there was no effect observed for *M. gaditana-* or *N. oculata*-infected larvae. **d** LPS exposure for 24 h after established symbiosis (24 h infection + 24h of incubation without free SSB01) does not influence SSB01 maintenance. **e** MyD88 inhibition enhances maintenance of SSB01 in larvae (*p*=0.0073, t=3.979), while *N. oculata* maintenance is not affected. All graphs show individual values plus mean ± SEM. ** indicates p<0.01 in two tailed paired (c,e) or unpaired (d) t-test.

Using our cell-type specific transcriptomic dataset, we compared the expression levels of immune-related genes (based on KEGG (Kanehisa and Goto, 2000) annotations for *E. pallida*) between the different cell-types (Fig. 2a, Fig. S2a). An array of immune genes was significantly down-regulated in symbiont-containing cells, compared to all other cell types (Fig. 2a). This down-regulation involved 10 KEGG pathways related to metazoan innate immunity with some genes present in multiple pathways, and multiple transcripts annotated as the same gene (e.g. TRAF3) (Fig. 2a), but it was the expression of genes associated with NF-κB and AP-1/Jun-related signalling pathways, such as toll-like receptor pathway (TLR, Fig. 2b) and tumour necrosis factor (TNF) pathways, that differed most drastically (∼20% of genes from each pathway) (Fig. 2a, Table S1). This demonstrates for the first time that symbiont phagocytosis massively suppresses transcription of immunity-related genes in the host at the cellular level. Moreover, in contrast to all previous organismal-wide analyses, we did not find the NF-κB transcription factor itself to be down-regulated in symbiotic cells (Mansfield et al., 2019; Wolfowicz et al., 2016; Mansfield et al., 2017) suggesting that its own transcriptional regulation is not required for symbiosis establishment. However, AP-1, a different transcription factor implicated in the TLR pathway is significantly downregulated in symbiont-containing cells (Fig. 2b).

Out of 382 unique genes involved in immunity, only a few are differentially regulated when comparing aposymbiotic cells from symbiotic and naïve larvae. Therefore, the transcriptional profile somewhat resembles the aposymbiotic state; however, some of the genes (11) are upregulated (Fig. 2a, Table S1). Thus, the explicit host cell response to symbiont phagocytosis is again distinct from the systemic response in the endoderm of infected *Aiptasia* larvae.

### Modulation of host innate immunity is involved in symbiosis establishment

The substantial suppression of host innate immunity (Fig. 2a) suggests that it is required for symbiosis establishment within the first 48 h after phagocytosis. To test this, we challenged aposymbiotic *Aiptasia* larvae with Lipopolysaccharides (LPS), a ligand for mammalian TLR commonly used to elicit an immune response, for 1 h before, and during infection with microalgae (Fig. 2c). Strikingly, LPS treatment massively decreased the proportion of symbiont-infected larvae to 28 % of the untreated control (Fig. 2c), reflecting previous observations in *Aiptasia* adult anemones (Detournay et al., 2012). We observed a similar, though less dramatic, effect for *C. velia*, but no significant effects for *N. oculata* and *M. gaditana* infected larvae (Fig. 2c). Surprisingly, once symbionts are intracellularly integrated for > 24 h, LPS treatment had no effect on symbiosis stability (Fig. 2d). Together, this suggests that for both the dinoflagellate symbiont and the apicomplexan-related ‘invader’, activation of host immunity pathways conflicts with the initiation of the symbiotic interaction. However, once symbionts are stably integrated into the host cell, symbiosis stability is not compromised by immune activation, likely due to the fact that a large proportion of host immune signalling is transcriptionally repressed by that time (Fig. 2a). The intracellular persistence time of *N. oculata* and *M. gaditana* microalgae is unaffected by LPS suggesting that their loss is independent of immune stimulation or occurs so quickly that any additional effect is undetectable.

In vertebrates, TLRs act as pattern recognition receptors (PRRs) and are involved in the elimination of pathogens; however, commensal bacteria have also been shown to exploit the TLR pathway by suppressing host immunity to promote host colonisation, suggesting that the host innate immune system responds distinctly to pathogens and symbionts (Franzenburg et al., 2012; Koch et al., 2018; Round et al., 2011). During colonisation, the central adapter protein MyD88 has emerged as a key player to manage the initiation of symbiotic pairings between host and beneficial microbe. MyD88 promotes re-establishment of bacterial homeostasis after antibiotic treatment in *Hydra vulgaris* and is required for an appropriate immune cell response during gut colonisation in zebrafish larvae (Franzenburg et al., 2012; Koch et al., 2018). Specifically, colonisation with beneficial microbial communities of zebrafish larvae induces a broad scale, *myd88*-dependent transcriptional reprogramming of immunity-related genes in the host, including suppression of MyD88 itself, as well as the transcription factor AP-1 (Koch et al., 2018), and in *H. vulgaris* specifically the AP-1 branch is effect during gut colonisation and in MyD88 silencing (Franzenburg et al., 2012). Strikingly, we also find that gene expression of large parts of the TLR pathway is significantly downregulated, including MyD88 (below detection threshold) and AP-1, after symbiont phagocytosis in *Aiptasia* larval host cells (Fig. 2a). To directly test whether decreased MyD88 activity affects retention of symbionts or *N. oculata* cells, we infected *Aiptasia* larvae with both for 24 h, and then inhibited the activity of MyD88 with an inhibitory peptide that blocks MyD88 homodimer formation (Loiarro et al., 2007). We found that MyD88 inhibition significantly increased the number of larvae that contained symbionts compared to the control; however, no such effect was observed for *N. oculata* (Fig. 2e). This suggests that inhibition of MyD88 facilitates symbiont retention within the first 48 h of infection. More broadly, suppression of MyD88 at the transcriptional level, and thus modulation of host innate immunity, may be a common strategy used in distinct symbiotic associations ranging from the vertebrate gut microbiome to cnidarian-dinoflagellate endosymbiosis to facilitate effective symbiont-host pairings.

### Only symbionts establish an intracellular niche, while all other microalgae are expelled

Phagocytosis is an ancient process by which cells internalise large particles from the environment. Originally utilised by single-celled organisms such as amoeba to acquire food, phagocytosis has evolved to become an important part of immunity to kill invading microbes in macrophages of higher metazoans (Flannagan et al., 2012). Accordingly, it seems likely that the non-symbiotic microalgae are cleared by intracellular digestion, characterised by the consecutive fusion of the nascent phagosome with endocytic vesicles to ultimately mature into the digestive phagolysosome. To our surprise, *N. oculata-* and *M. gaditana*-containing phagosomes appeared to be void of the lysosomal-associated membrane protein 1 (LAMP1), a marker protein typically associated with late endo- and lysosomes destined for digestion. In contrast, LAMP1 was found to be weakly associated with *C. velia*-containing phagosomes, and heavily decorating symbiosomes, the organelle in which the symbiont resides, shortly after uptake (Fig. 3a, Videos S1-3). Thus, it is the symbionts that rapidly establish a LAMP1-positive niche in which they can persist intracellularly, and not the non-symbiotic ‘invaders.’ This then raises the question: how are non-symbiotic microalgae, destined for removal (Fig. 1d), cleared by the host cell after phagocytosis?

**Figure 3.**
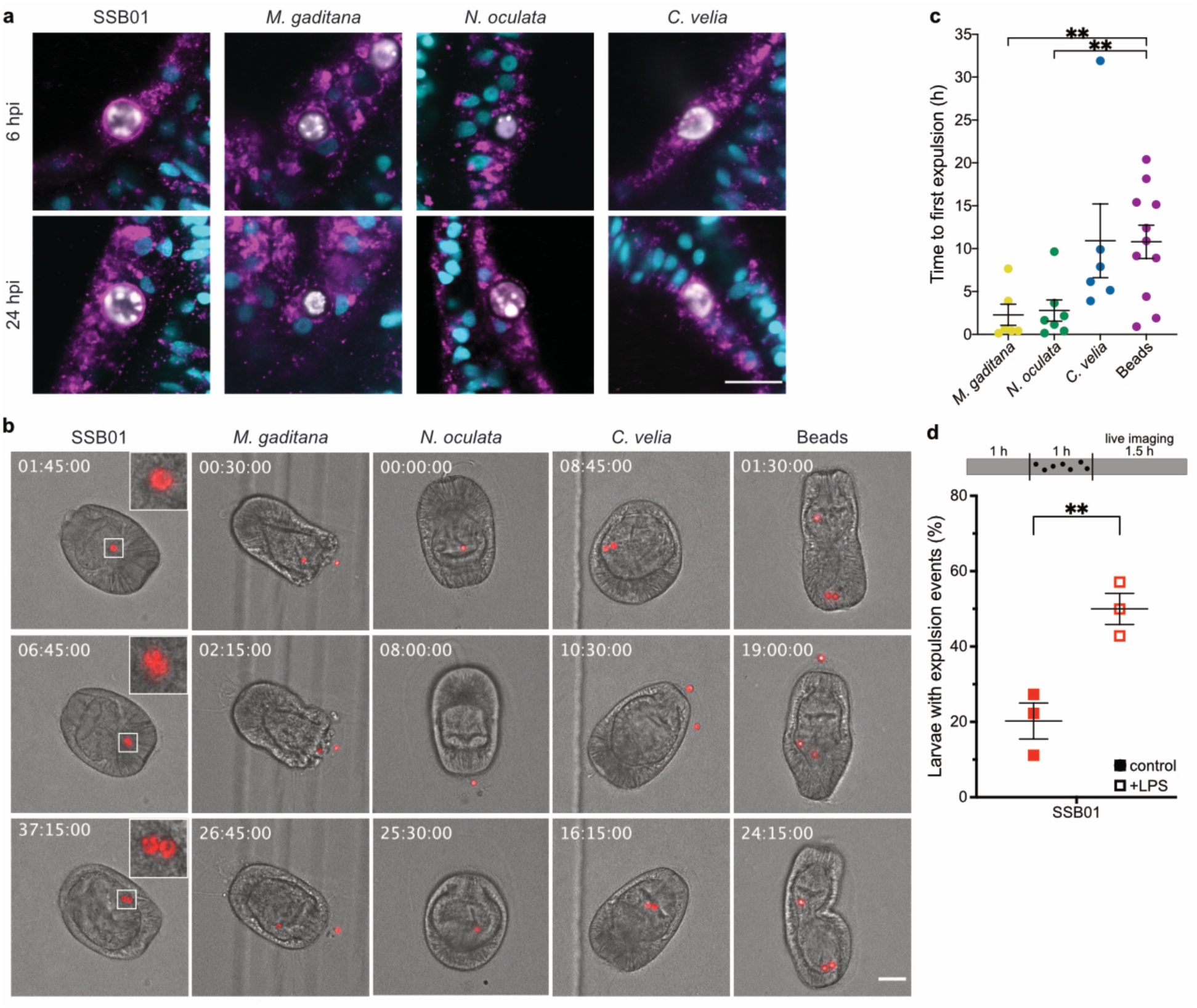
Non-symbiotic microalgae are removed by expulsion. **a** Only SSB01 is maintained in a LAMP1-positive symbiosome, as early as 6 hpi, while the other microalgae show no, or only slight, LAMP1 staining. LAMP1 (magenta); Hoechst (cyan); microalgae (white). Scale bar represents 10 µm. **b** During live imaging of larvae, SSB01 (symbiont) did not appear to be expelled and were also seen to replicate within the host endoderm. *M. gaditana*, *N. oculata*, and *C. velia* and beads were all expelled at some point during imaging. Timestamps indicate time after the start of imaging (hh:mm:ss), which coincides with 24 hours post infection. Stills are DIC and red autofluorescence of algal photosynthetic pigments, scale bar represents 30 µm, inserts are zoomed 3-fold. For corresponding videos see videos S6-10. **c** Time until first expulsion after onset of imaging is shorter with *M. gaditana* (*p*=0.0084, t=3.03, compared to beads) and *N. oculata* (*p*=0.008, t=3.031, compared to beads) than with *C. velia* or latex beads (only time for expelled particles shown). **d** Reduced amounts of infected larvae upon LPS treatment is due to higher expulsion rate. LPS exposure for 1h before and during 1h of infection significantly enhances number of larvae with expulsion events as observed within 1.5h of live imaging (*p*=0.0091, t=4.724). In all graphs, lines indicated mean ± SEM, ** indicated *p*<0.01 in two-tailed unpaired t-test.

To monitor elimination of non-symbiotic microalgae, we established live imaging to observe their fate over time. After 24 hpi, larvae were embedded in low-gelling agarose (LGA) and imaged every 15 minutes for 48 h. We selected larvae that appeared to have intracellular microalgae, indicated by continuous and synchronous movement of both the algal cell and the rotating larva (Video S4); this was clearly distinct from the non-synchronous movement of microalgae located in the gastric cavity (* in Video S4). Intracellular symbionts were maintained inside *Aiptasia* larvae for the entire duration of the observation period (Fig. 3b, Video S5, Table S2), with the exception of one larva at one time point in which a pair of symbionts was seen moving asynchronously in the gastric cavity (Video S6). Furthermore, symbionts frequently replicated (11 replication events/13 larvae) suggesting that the immobilisation in LGA does not affect larval or symbiont physiology, nor symbiosis stability.

To our surprise, most larvae that had phagocytosed *M. gaditana* cells (4/7)*, N. oculata* cells (7/8), and *C. velia* cells (6/8) expelled the microalgae rather than digesting them (Fig. 3b, Video S7-9, Table S2). After expulsion, the microalgae appeared intact and healthy based on the retention of their autofluorescence and their ability to re-infect larvae. Re-acquisition of non-symbiotic microalgae by larvae occurred frequently (Table S2). To test whether this was a common response to inert particles in addition to non-symbiotic microalgae, we imaged larvae containing intracellular polystyrene beads of the same size as the microalgae. We saw that all of the beads were also both expelled (11/11) and reacquired (Table S2, Video S10). Interestingly, we found that initial expulsion of *C. velia* cells and beads occurred later during imaging when compared to the expulsion of *N. oculata* and *M. gaditana* cells (Fig. 3c); in one event, a *C. velia* algal cell replicated during imaging (Video S9). Together, this indicates that non-lytic expulsion, and not phagolysosomal digestion, is a general response to the uptake of non-symbiotic particles in *Aiptasia* larvae. Uptake and subsequent removal appear to occur non-specifically and continuously, as a trial-and-error mechanism, to probe for suitable particles (e.g. symbionts) in the environment.

### Non-lytic expulsion (vomocytosis) is negatively regulated by ERK5

Interestingly, in both phagotrophic amoeba and vertebrate macrophages, the fungus *Cryptococcus neoformans* and certain *Candida* species are known to be expelled via a process referred to as vomocytosis (Bojarczuk et al., 2016; Smith and May, 2013). The host phagocyte remains intact, thus preventing immune stimulation, which is thought to promote dissemination of the pathogen (Alvarez and Casadevall, 2006; Ma et al., 2006). To directly test if immune activation via LPS treatment induces symbiont loss by expulsion rather than reduced uptake, we used our live-imaging assay to monitor their fate. We found that in the presence of LPS, symbionts are much more likely to be expelled (Fig. 3d). Very early after infection (~1-6 hpi), we observed a baseline expulsion of symbionts that we did not observe in earlier experiments (Fig 3d), likely due to the fact that earlier experiments were monitoring a more stable, well-established symbiosis (24 hpi). Thus, immune stimulation by LPS counteracts the robustness of symbiosis establishment due to the increase in expulsion events of newly phagocytosed symbionts.

In vertebrate macrophages, vomocytosis is negatively regulated by MAPK 7 (ERK5), as well as the MAPK kinase, MEK5. ERK5 inhibition by XMD17-109 significantly increased rates of vomocytosis (Gilbert et al., 2017). *Aiptasia* contains several MAP kinases and one clear ERK5 homolog, with a conserved ATP binding site (aa 61-69 in *H. sapiens* ERK5 – the target of XMD17-109), as well as a clear MEK5 homolog (Fig. S3). To examine whether the function of ERK5 as a negative regulator of vomocytosis is conserved in *Aiptasia*, we incubated *Aiptasia* larvae shortly before, and during, a 24 h infection period with the inhibitor. Intriguingly, we found that the number of larvae infected with symbionts was massively reduced upon ERK5 inhibition (Fig. 4a). Moreover, upon inhibition of ERK5, LAMP1 accumulation on the symbiosome does not occur, resembling non-symbiotic microalgae-containing phagosomes (Fig. 4b + c). This suggests that ERK5 inhibition increases symbiont expulsion via vomocytosis and impairs the niche formation that is required to initiate a stable partnership between endosymbiont and host cell. More broadly, vomocytosis is an evolutionarily conserved mechanism utilised by amoeba, cnidarians, and vertebrates as a means to clear unwanted particles from phagocytic cells (Alvarez and Casadevall, 2006; Ma et al., 2006; Watkins et al., 2018, this study).

**Figure 4.**
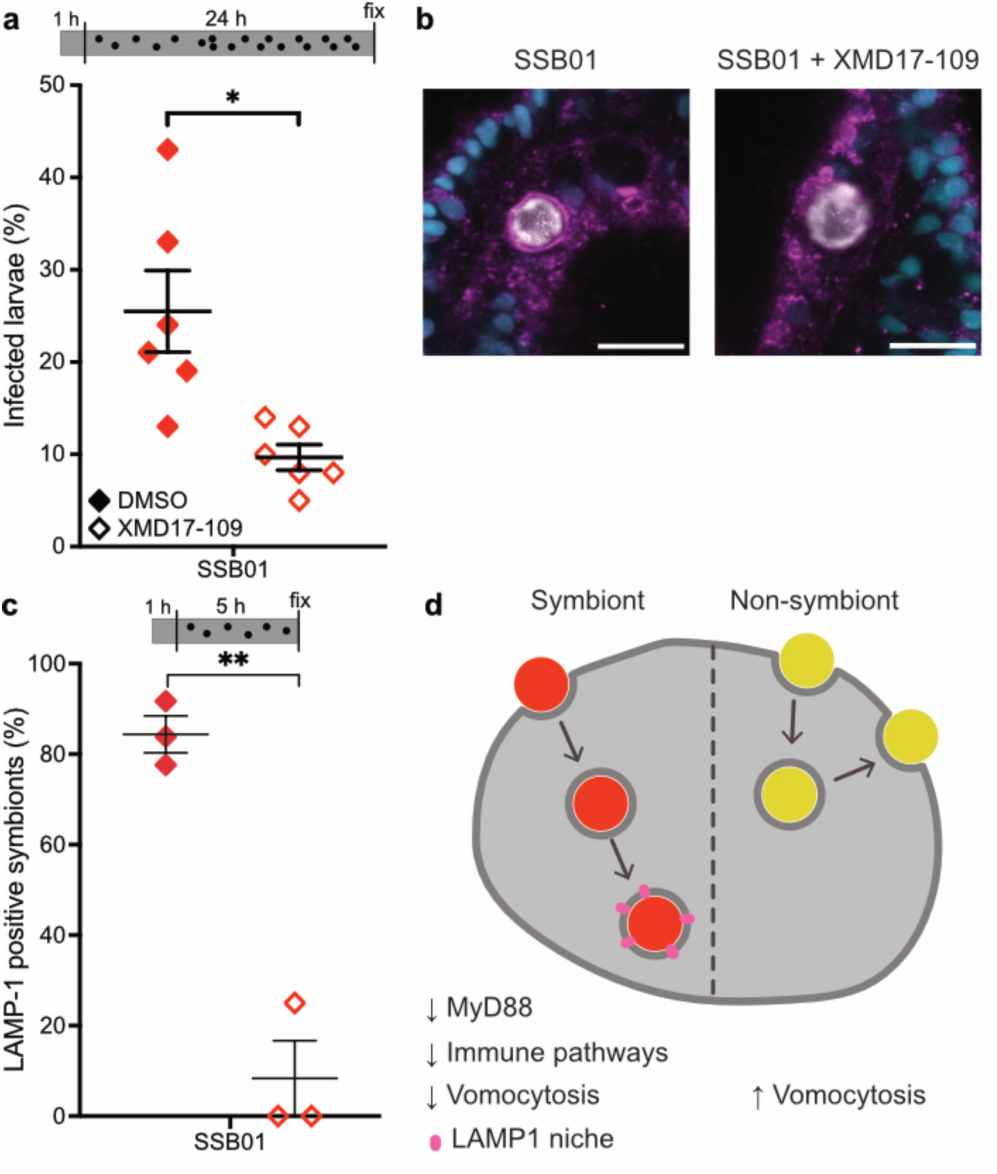
ERK5 is a negative regulator of vomocytosis. **a** ERK5 inhibition with XMD17-109 reduces fraction of symbiotic larvae, when pre-treated for 1h and treated during 24h infection. Lines indicate mean ± SEM, * indicates *p*=0.0135, t=3.729 in two-tailed paired t-test. **b L**arvae with normal ERK5 activity are able to form a LAMP1-positive symbiosome, while larvae treated with XMD17-109 to inhibit activity of ERK5 show massively reduced accumulation of LAMP1 when pre-treated for 1h and treated during 5 h infection. LAMP1 (magenta); Hoechst (cyan); microalgae (white). Scale bar represents 10 µm. **c** ERK5 inhibition significantly reduces fraction of symbionts with LAMP1 accumulation. Lines indicate mean ± SEM, ** indicates *p*=0.0043, t=15.14 in two-tailed paired t-test. **d** Model of algal uptake in *Aiptasia* endodermal cells. While uptake of Symbionts (SSB01, red) leads to downregulation of immunity genes, until a functional symbiosomes is formed, where this is no longer necessary, other non-symbiotic microalgae (yellow) elicit no strong transcriptomic response in the host cells and are expelled by a vomocytosis-like process.

### Conclusions

We propose a model in which aposymbiotic *Aiptasia* larvae constitutively acquire particles from the environment. By default, phagocytosed, non-symbiotic microalgae are removed by vomocytosis. Acquisition of true symbionts leads to suppression of host innate immunity pathways, likely the AP-1 branch of the highly conserved TLR signalling, by targeting MyD88. Modulation of host innate immunity is crucial for symbionts to escape expulsion by vomocytosis and promote intracellular niche formation (Fig. 4c). More broadly, the initiation of the endosymbiotic association between dinoflagellate symbionts and cnidarian host cells depends on subverting the innate immune system; specifically, the removal of invaders by expulsion, and not by phagolysosomal digestion.

Phagocytosis is an ancient mechanism used by unicellular organisms to acquire and digest food (Hartenstein and Martinez, 2019). Microbial killing by professional macrophages in higher metazoans is thought to derive from phagotrophy (Boulais et al., 2010; Hartenstein and Martinez, 2019). Interestingly, the core TLR/NF-κB pathway has recently been identified in choanoflagellates suggesting that unicellular organisms may have used phagocytosis for both prey sensing and immunity (Richter et al., 2018). Although the evolutionary trajectory of phagocytosis is unclear, it seemed likely that intracellular digestion is a primitive and highly conserved defence mechanism. Thus, it was highly unexpected that symbionts have to escape vomocytosis to efficiently establish a stable intracellular symbiotic relationship.

In this context, it is noteworthy that unicellular, phagotrophic amoeba have two redundant, yet mechanistically distinct expulsion mechanisms: constitutive exocytosis and vomocytosis (Watkins et al., 2018). Constitutive exocytosis results in the expulsion of undigested remnants of food; alternatively, pathogens such as *C. neoformans*, which are protected from digestion by their polysaccharide capsule (Lee et al., 1995), activate a second route of expulsion, vomocytosis, possibly in an effort to avoid infection or halt the acquisition of new food. Expulsion of indigestible material, non-symbiotic microalgae, or even pathogenic microbes may be effective for small, motile organisms that are in constant contact with the environment, but animals with complex body plans and organ systems are more susceptible to lasting infections by expelled, yet internally trapped, microorganisms. Therefore, vomocytosis of unwanted and indigestible particles may be an evolutionarily ancient defence mechanism that required stricter regulation in higher metazoans, which focus on microbial killing. This may explain why this phenomenon has been overlooked in multicellular organisms until recently (Alvarez and Casadevall, 2006; Ma et al., 2006).

Cnidarians, such as corals and anemones, are evolutionarily ancient animals with simple body plans, yet possess cells with complex immune capacities. Here, we show that the endodermal cells of cnidarian larvae use distinct mechanisms of innate immunity to abet a post-phagocytic trial-and-error mechanism for selection of suitable symbionts from the environment. This study provides valuable insight into regulation of host immunity by symbionts and demonstrates that the primitive and versatile functions of phagocytosis are exploited by these basal organisms to support symbiosis and host health.

## Acknowledgements

We thank Dinko Pavlinic and Vladimir Benes (Genecore Facility, EMBL Heidelberg) for assistance with the SmartSeq2 protocol and sequencing library preparation; David Ibberson (Deepseqlab, Heidelberg University) for assistance with the SmartSeq2 protocol; Carsten Rippe for access to the BioAnalyzer; Liz Hambleton and Ira Maegele for help with antibody purification; Bruno Gideon Bergheim for initiating live imaging of *Aiptasia* larvae; Friedrich Frischknecht, Robin May, Thomas Holstein and Steffen Lemke for advice and comments; Robin May for comments on the manuscript.

Funding was provided by the Deutsche Forschungsgemeinschaft (DFG) (Emmy Noether Program Grant GU 1128/3–1) and the H2020 European Research Council (ERC Consolidator Grant 724715) to AG, a scholarship to SR by the CellNetworks Excellence Cluster (Heidelberg University) Postdoctoral Program, and a PhD scholarship within the Graduate School “Evolutionary Novelty & Adaptation by the Baden-Württemberg Landesgraduiertenförderung Program to PAV.

## Author Contributions

Conceptualization, M.R.J., S.R. and A.G.; Methodology, M.R.J., S.R., PAV and A.G.; Software, P.A.V. and S.G.G.; Formal Analysis, M.R.J., S.R., Investigation, M.R.J., S.R.; Resources, A.G.; Data Curation, P.A.V. and S.G.G.; Writing - Original Draft, M.R.J., S.R. and A.G., Writing –Review & Editing, M.R.J., S.R. and A.G.; Visualization, M.R.J., S.R.; Supervision, A.G., Project Administration, M.R.J., S.R. and A.G.; Funding Acquisition, A.G.

## Supplementary Information

is available for this paper.

Correspondence and requests for materials should be addressed to A.G.

## Methods

### Live Organism Culture and Maintenance

#### Algal maintenance

For infection experiments of *Aiptasia* larvae, we used *Breviolum minutum* clade B (SSB01) (Xiang et al., 2013), *Microchloropsis gaditana* CCMP526 (NCMA, Bigelow Laboratory for Ocean Sciences, Maine, USA), *Nannochloropsis oculata*, and *Chromera velia* (NORCCA K-1276). All cultures were grown in cell culture flasks in 0.22 µm filter-sterilized 1X Diago IMK medium (Wako Pure Chemicals, Osaka, Japan) on a 12-h light:12-h dark (12L:12D) cycle under 20–25 µ mol m−2 s−1 of photosynthetically active radiation (PAR). Sterile stock cultures of SSB01, *N. oculata*, and *C. velia* were grown at 26 ºC and *M. gaditana* at 18 ºC. All algal cultures to be used for infections were kept at 26 ºC for a minimum of 3 days prior to infection.

#### Aiptasia spawning and larval culture conditions

*Aiptasia* clonal lines F003 and CC7 (Carolina Biological Supply Company #162865; Burlington, USA) were induced to spawn following the previously described protocol (Grawunder et al., 2015). *Aiptasia* larvae were maintained in glass beakers in filter-sterilized artificial sea water (FASW) at 26 ºC and exposed to a 12L:12D cycle.

### Infection assay

Three biological replicates (e.g. distinct spawning events) of naturally aposymbiotic *Aiptasia* larvae were collected and diluted to a concentration of 300-500 larvae per millilitre of FASW in glass beakers. Between 4- and 8-days post fertilisation (dpf), the larvae were infected with 1.0 × 10^5^ algal cells/ml of the respective algal types. Beakers were kept at 26 ºC and exposed to a 12L:12D cycle. After a 24-h infection, the larvae were washed to remove the microalgae and fresh FASW was added.

#### Quantification of infection efficiency

Infected larvae were fixed at 1-, 2-, 3-, 6-, and 10-day(s) post infection (dpi) using 4% formaldehyde solution (#F1635, Sigma-Aldrich) for 30 minutes at room temperature (RT), followed by two washes in 0.1% Triton X-100 (PBS-Triton) (#3051, Carl Roth GmbH), and mounted in 87% glycerol (#G5516, Sigma-Aldrich) in PBS with the addition of 2.5 mg/ml 1,4-Diazabicyclo[2.2.2]octan (DABCO) (#D27802, Sigma-Aldrich). At least 50 larvae per replicate per algal type were counted, and representative differential interference contrast (DIC) and epi-fluorescent images of the algal autofluorescence were taken. Microscopic analysis was conducted with a Nikon Eclipse Ti inverted microscope using a Nikon Plan Fluor 40x air objective. Images were processed with Fiji software (Schindelin et al., 2012).

### Imaging and staining procedures in *Aiptasia* larvae

#### Fluorescent staining for f-actin

Infected larvae were fixed 24 hpi using 4% formaldehyde solution for 30 minutes at RT, followed by one wash in 0.05% Tween20 (#P7949, Sigma-Aldrich) in PBS (PBS-Tween20) for 5 minutes. For permeabilisation, larvae were rotated in 1.5 ml Eppendorf tubes at 0.25 rpm in a solution of 1% PBS-Triton and 20% DMSO (#67-68-5, fisherscientific) for 1 h at RT. For blocking, the permeabilisation solution was exchanged with 5% normal goat serum (#005-000-121, Jackson ImmunoResearch Laboratories, Inc.) in 0.05% PBS-Tween20. Larvae remained in the blocking buffer for 30 minutes at RT while rotating. After two washes in 0.05% PBS-Tween20, larvae were incubated in Phalloidin Atto 565 (#94072, Sigma-Aldrich) diluted 1:200 in 0.05% PBS-Tween20 on a rotor and protected from light. Larvae were washed three times in 0.05% PBS-Tween20 before incubation in 10 µg/ml Hoechst (#B2883, Sigma-Aldrich) diluted in Tris-buffered saline, pH 7.4; 0.1% Triton X-100; 2% bovine serum albumin (#A7906, Sigma-Aldrich); 0.1% sodium azide (#S2002, Sigma-Aldrich) for 30 minutes on a rotor at RT in the dark. Larvae were washed three times for 5 minutes with 0.05% PBS-Tween20 and washed step-wise into glycerol from 30% to 50% to 100% and finally mounted. Confocal microscopic analysis was carried out with a Leica TCS SP8 stand using 63x glycerol immersion objective (NA 1.30), Leica LAS X software and Fiji software (Schindelin et al., 2012). Hoechst, Atto 565, and symbiont autofluorescence were excited with 405, 561, and 633 nm laser lines, respectively. Fluorescence emission was detected at 410-501 nm for Hoechst, 542-641 nm for Phalloidin Atto 565, and 645-741 for symbiont autofluorescence.

#### Live imaging

For live imaging of infected larvae, chambers were prepared by placing two 2.5 mm x 5 mm strips of non-toxic double-sided tape (TES5338, Tesa) at the periphery of 35 mm µ-Dishes (# 81166, Ibidi). 1.5% low gelling agarose (LGA) (#A4018, Sigma-Aldrich) was dissolved in FASW by heating to 80 ºC until liquid. The liquid LGA was cooled to and kept at 37 ºC until use. 300 larvae in a volume of 1 ml were placed in a 1.5 ml Eppendorf centrifuge tube and quickly vortexed using a Sprout mini centrifuge (#552021, Biozym Scientific GmbH) to pellet larvae. The larvae and 1.5% LGA at 37 ºC were mixed for a final LGA concentration of 1.14% LGA. The mixture was pipetted into the center of the prepared imaging chamber and a glass coverslip was pressed on top of the droplet as to adhere on either end to the double-sided tape. The Ibidi plate was filled with 2 ml of FASW. Microscopic analysis was conducted with a Nikon Eclipse Ti inverted microscope using a Nikon Plan Fluor 20x air objective, with images taken every 15 minutes in DIC and TexasRed channels. Images were processed with Fiji software (Schindelin et al., 2012).

#### Immunofluorescence staining (LAMP1)

Larvae were fixed for 45 minutes in 4% formaldehyde at RT, followed by three washes in 0.2% PBS-Triton and one wash in PBS. Larvae were then permeabilised in PBS-Triton for 1.5 h at RT, followed by blocking in 5% normal goat serum and 1% BSA in PBS-Triton for 1 h. Primary antibody (rabbit-α-LAMP1) (Voss et al., 2019) was diluted 1:100 in blocking buffer and incubated overnight at 4°C. After three washes in PBS-Triton, the secondary antibody (goat-α-rabbit AlexaFluor 488, #ab150089, abcam) was diluted 1:500 in blocking buffer and incubated for 1.5 h at RT. Larvae were washed two times in PBS-Triton, followed by a 15-minute incubation with 10 µg/ml Hoechst protected from light at RT, and two final washes in PBS-Triton and one in PBS before mounting in 87% glycerol. Larvae were imaged on a Leica TCS SP8 confocal laser scanning microscope using a 63x glycerol immersion lens (NA 1.30) and Leica LAS X software. Hoechst, Alexa 488, and symbiont autofluorescence were excited with 405, 496, and 633 nm laser lines, respectively. Fluorescence emission was detected at 410-501 nm for Hoechst, 501-541 nm goat-α-rabbit AlexaFluor 488, and 645-741 for symbiont autofluorescence.

### Transcriptome

#### Computational methods

RNA sequencing data from Voss, et al. (2019) was complemented with data for larvae infected with *M. gaditana*, acquired in the same way. Paired reads were mapped to the *Aiptasia* genome version GCF_001417965.1 using HISAT2 version 2.1.0 at default settings, except –X 2000 --no-discordant --no-unal --no-mixed. Transcripts were quantified in Trinity v2.5.1 using salmon v0.10.2 at default settings. Principal component analysis was conducted using a perl script supplied with Trinity for all samples. Differential expression was analysed using in R! v3.5.2 (R Core Team, 2018). DEseq2 (Love et al., 2014) (log2-fold change ≥ 1, adjusted p-value ≤ 0.05). Graphing of results was restricted to pathways involved in innate immunity as found in KEGG (Kanehisa and Goto, 2000), using the R! package “ComplexHeatmap” (Gu et al., 2016) in KNIME (Berthold et al., 2007). The custom KNIME workflow used for processing and analysing the data are freely available from the corresponding author upon request.

### Exogenous immune perturbations

#### LPS/ ERK5 inhibitor treatment

*Aiptasia* larvae were washed and diluted to a concentration of 300-500 larvae/ml and were then incubated with 20 µg/µl LPS (lipopolysaccharides from *Escherichia coli* O127:B8, Sigma-Aldrich), 1 µm XMD17-109 (0.1% DMSO [Dimethylsulfoxid, Carl-Roth]), 0.1 % DMSO, or without for 1 hour. Algal cultures were then added to a final concentration of 1 × 10^5^ microalgae/ml of the respective algal types and incubated at 26 ºC and exposed to a 12L:12D cycle. After a 24-hour infection, the larvae were fixed in 4% formaldehyde for 30 minutes, washed in PBS, and mounted in 100% glycerol for counting. Infection status was quantified in at least 100 larvae per infection using a Nikon Eclipse Ti epifluorescent microscope, utilising algal autofluorescence.

#### LPS post-infection treatment

Washed and diluted *Aiptasia* larvae (300-500 / ml) were mixed with *B. minutum* (1×10^5^ microalgae / ml) and left for infection at 26 °C with a 12L:12D cycle for 24 hours. They were then washed/filtered to remove excess microalgae and incubated further for 24 hours. Half of the infected larvae were exposed to 20 µg/ml of LPS for 24 hours, before fixation with 4% formaldehyde. They were then washed in PBS and mounted in glycerol. Infection status was assessed in at least 100 larvae per infection using a Nikon Eclipse Ti epifluorescent microscope, utilising algal autofluorescence.

#### Early infection LPS treatment live imaging

*Aiptasia* larvae (300-500) were either incubated with 20 µg/ml LPS or not for 1 hour, before infection with *B. minutum* (1×10^5^ microalgae / ml) for 1 hour. They were then mounted in µ-Dishes as described previously and then imaged on a Nikon Eclipse Ti inverted microscope using a Nikon Plan Fluor 20x air objective, with images taken every 5 min in DIC and TexasRed channels. Images were processed with Fiji software (Schindelin et al., 2012). For each independent experiment (n=3), between 7 and 14 larvae were observed.

#### MyD88 inhibitor peptide treatment

6 dpf larvae were infected with 1.0 × 10^5^ cells/ml of SSB01 or *N. oculata* for 24 hours. Infected larvae were washed with FASW to remove microalgae from water and 10 units penicillin and 10 µg streptomycin (Sigma-Aldrich) were added. Infected larvae were incubated for 24 hours at 26 ºC under a 12L:12D cycle with either 50 µM of MyD88 inhibitor peptide or 50 µM control peptide (NBP2-29328, Novus Biologicals). Larvae were fixed in 4% formaldehyde and infection efficiency was quantified as previously described.

#### LAMP1 assessment after ERK5 inhibitor treatment

*Aiptasia* larvae were exposed to 1 µM ERK5 inhibitor (XMD17-109) or not for 1 h prior to and during infection with 1.0 × 10^5^ cells/ml of SSB01 for 5 h. Larvae where then stained for LAMP1 as described above. Fracture of symbionts with strong LAMP1 staining were assessed on a Leica TCS SP8 confocal laser scanning microscope using a 63x glycerol immersion lens (NA 1.30) and Leica LAS X software. For control 20 larva each were evaluated, due to low infection numbers in XMD17-109 treated larvae, only 8, 4 and 7 larvae could be evaluated in the replicates.

#### ERK5 / MEK5 phylogeny

To resolve the evolutionary history and classification of *Aiptasia* MAPKs and MAP2Ks we first used well-defined ‘bait’ sequences (human MAPK7 and MAP2K5) and reciprocal BLAST searches to identify and collate the MAPK and MAP2K repertoire in *Aiptasia. Aiptasia* sequences were then used to retrieve other MAPK and MAP2K sequences of animals including key cnidarian species from public databases using BLASTP. *H. echinata* sequences were downloaded from https://research.nhgri.nih.gov/hydractinia/download/ and manually curated and translated. Resulting protein sequences were deduplicated and then aligned using ClustalW (GONNET, goc: 3, gec: 1.8). Then automated trimming was performed using trimAI (using standard parameters; http://trimal.cgenomics.org/, (Capella-Gutierrez et al., 2009)). Best-fitting amino acid substitution models were determined using ModelFinder (-m MF -msub nuclear -nt AUTO) within iqTree and Maximum likelihood trees generated. Trees were finalised using FigTree 1.4.4 (http://tree.bio.ed.ac.uk/software/figtree/) and Adobe Illustrator CC 2018. Original tree files are available upon request

#### Statistical notes

Statistic test and number of replicates are given in text or figure legends. In general, two-sided t-test were performed using GraphPad Prism version 8.2 for Mac, GraphPad Software, La Jolla California USA, www.graphpad.com. Paired t-test were performed for all experiments, except for experiments represented in Fig. 3 where experimental set-up did not permit side-by-side execution and unpaired t-test were used instead. Individual experiments from distinct samples (n) are shown as data points.

#### Data Availability

The datasets generated during and/or analysed during the current study are available upon request.

#### Code Availability

All custom codes are available upon request from the corresponding author.

## Supplemental Information

**Fig. S1.**
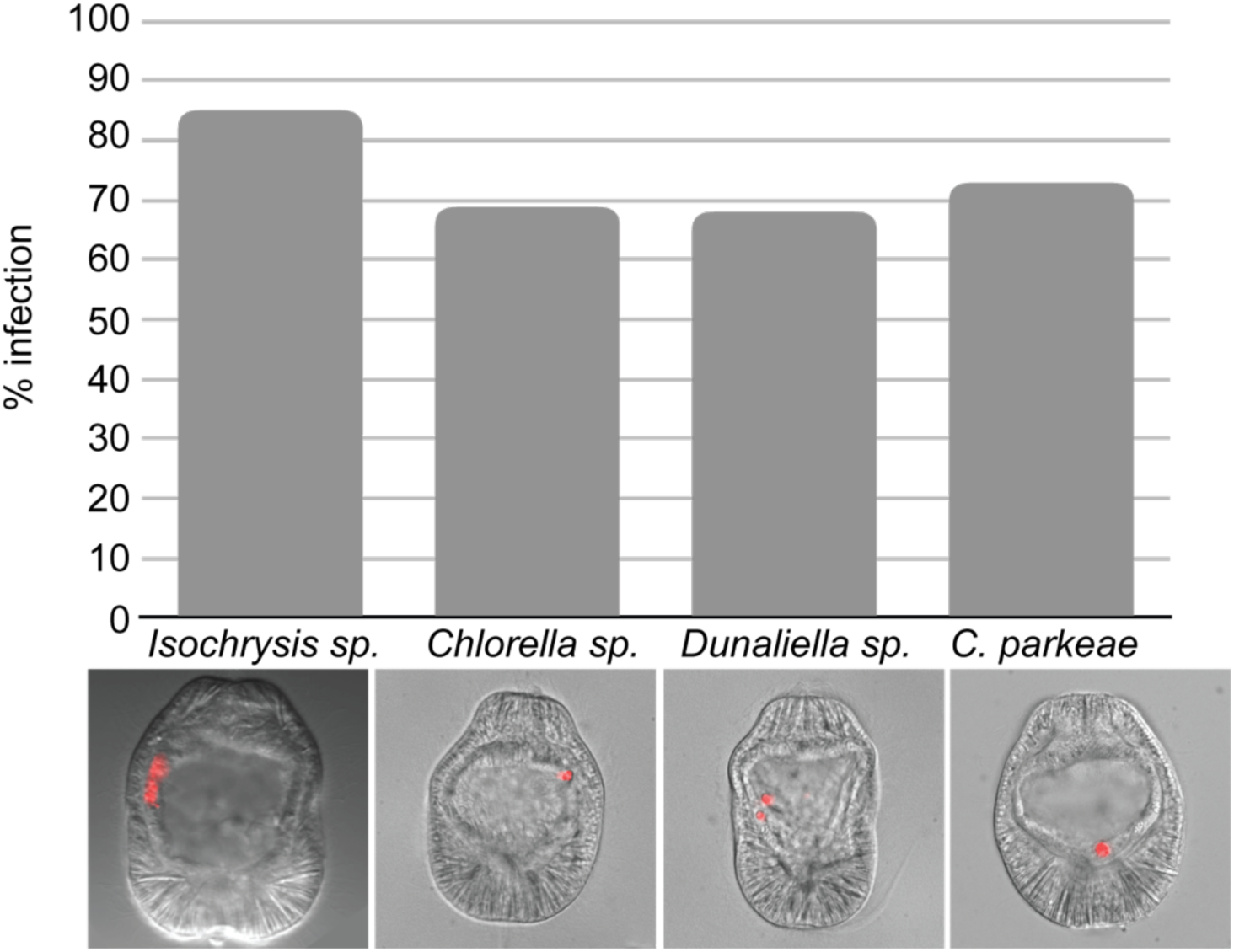
Additional microalgae screened: *Isochrysis sp., Chlorella sp., D. salina, and C. parkaea. Aiptasia* larvae were infected at 4-6 days post fertilisation (dpf) for 24 hours and were washed into fresh FASW. Images are DIC and red autofluorescence of algal photosynthetic pigments.

**Fig. S2.**
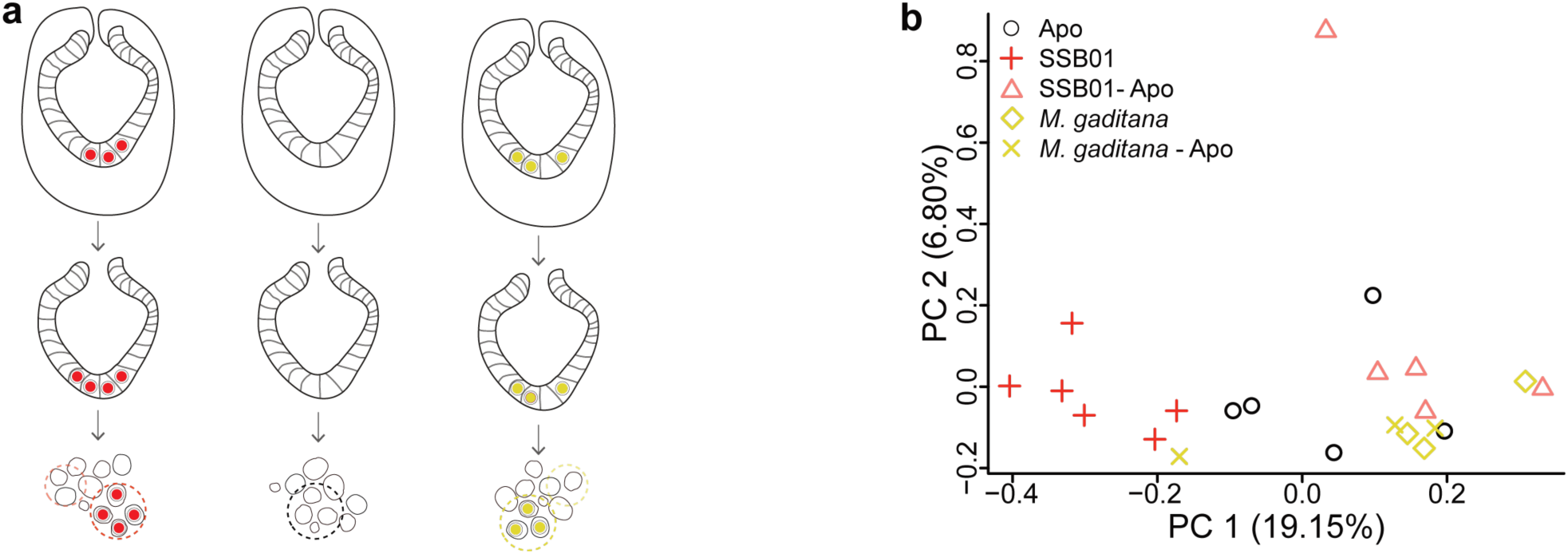
**a** Schematic of *Aiptasia* larvae used for cell-type specific sequencing. Ectodermal cells were removed resulting in only endodermal cells, which were dissociated and selected for based on contents: aposymbiotic cells from symbiotic larvae (SSB01-Apo), symbiotic cells from symbiotic larvae (SSB01), aposymbiotic cells from aposymbiotic larvae (Apo), cells containing *M. gaditana* from larvae infected with *M. gaditana* (Mg), and aposymbiotic cells from larvae infected with *M. gaditana* (Mg-Apo). **b** Principal Component Analysis (PCA) plot of host gene expression in different conditions.

**Fig. S3.**
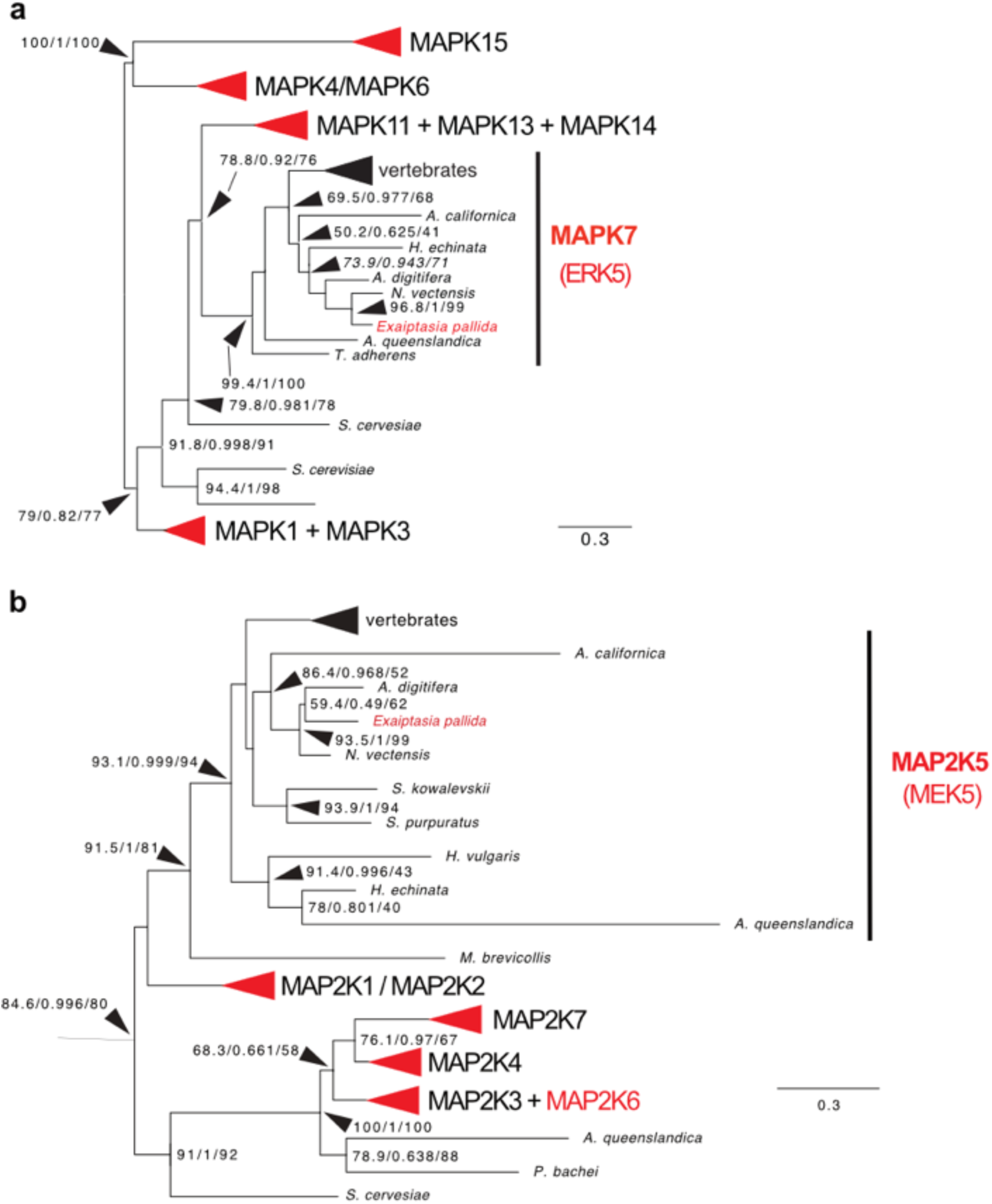
Phylogenetic analysis of ERK5 and MEK5 from *Aiptasia*. **a** + **b** are collapsed trees of Aiptasia MAPK (a) or MAP2K (b) in comparison to several other cnidarian and vertebrate species. Red arrowheads or writing indicate presence of an *Aiptasia* homolog. Both *Aiptasia* ERK5 and MEK5 cluster within ERK5 (MAPK7) or MEK5 (MAP2K5), respectively. Full tree can be accessed through file S1 and S2.

**Table S1.**
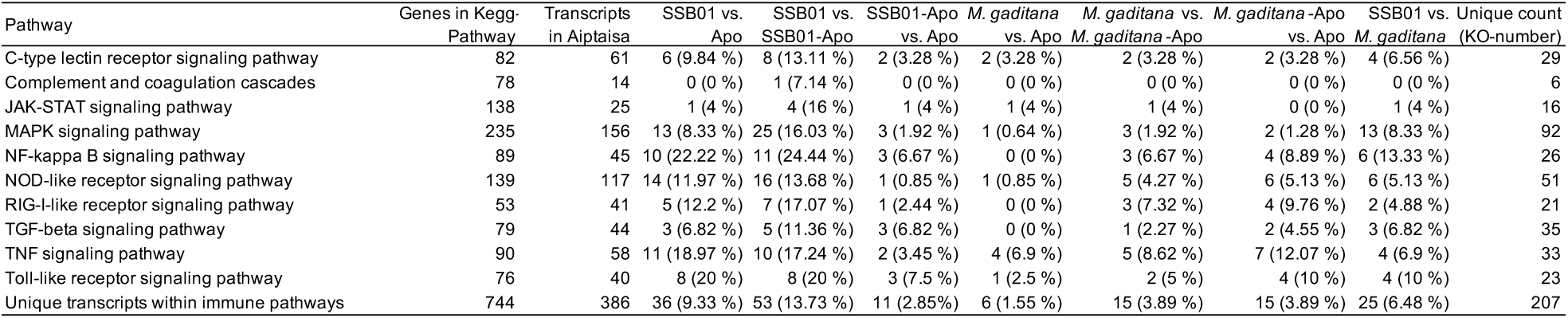
Statistics of transcriptional suppression of host cell immunity Enumeration of modulation of innate immunity genes over 10 immune pathways, with some genes present in multiple pathways, and multiple transcripts annotated as the same gene (e.g. TRAF3)

**Table S2.**
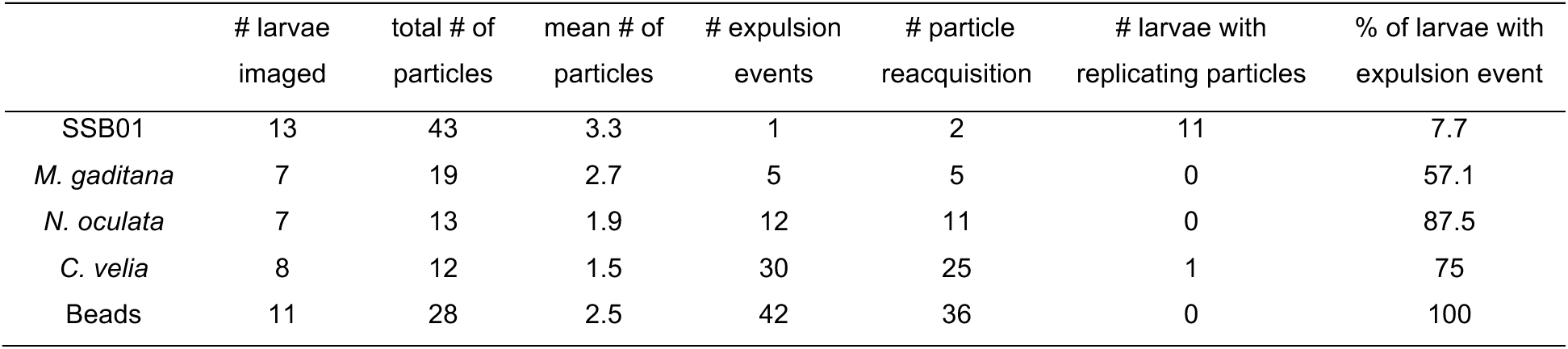
Live imaging statistics

### Video S1

3D reconstruction of LAMP1 staining in *Aiptasia* larvae infected with SSB01. LAMP1 is stained in magenta, DNA is stained with Hoechst in cyan, and autofluorescence of symbiont is shown in white.

### Video S2

3D reconstruction of LAMP1 staining in *Aiptasia* larvae infected with *N. oculata*. LAMP1 is stained in magenta, DNA is stained with Hoechst in cyan, and autofluorescence of symbiont is shown in white.

### Video S3

3D reconstruction of LAMP1 staining in *Aiptasia* larvae infected with *C. velia*. LAMP1 is stained in magenta, DNA is stained with Hoechst in cyan, and autofluorescence of symbiont is shown in white.

### Video S4

Intracellular / attached microalgae (SSB01) move in cohesion with the larva, whereas non-intracellular microalgae (*) clearly move independently within the gastric cavity. Autofluorescence of microalgae is shown in red. Timestamp is given in hh:mm:ss.

### Video S5

Long term imaging of larva infected with SSB01 (symbiont). SSB01 can be seen dividing at around 8 h after start of imaging. Auto fluorescence of microalgae is shown in red. Timestamp is given in hh:mm:ss.

### Video S6

Z-stack of SSB01 within the gastric cavity for one time point during acquisition. Auto fluorescence of microalgae is shown in red. Timestamp is given in hh:mm:ss.

### Video S7

Long term imaging of larva infected with *M. gaditana*. *M. gaditana* can be seen being expelled and taken up again. Auto fluorescence of microalgae is shown in red. Timestamp is given in hh:mm:ss.

### Video S8

Long term imaging of larva infected with *N. oculata*. *N. oculata* can be seen being expelled and taken up again. Auto fluorescence of microalgae is shown in red. Timestamp is given in hh:mm:ss.

### Video S9

Long term imaging of larva infected with *C. velia*. *C. velia* can be seen being expelled and taken up again. Auto fluorescence of microalgae is shown in red. Timestamp is given in hh:mm:ss.

### Video S10

Long term imaging of larva infected with Beads. Beads can be seen being expelled and taken up again. Auto fluorescence of microalgae is shown in red. Timestamp is given in hh:mm:ss.

### Supplementary File 1

Phylogenetic tree of MAPK, with focus on ERK5 (MAPK7).

### Supplementary File 2

Phylogenetic tree of MAP2K, with focus on MAP2K5 (MEK5).

