## Supplemental Figures and Tables for "Dinoflagellate symbionts escape vomocytosis by host cell immune suppression"

**Supplemental Information**

**Fig. S1**

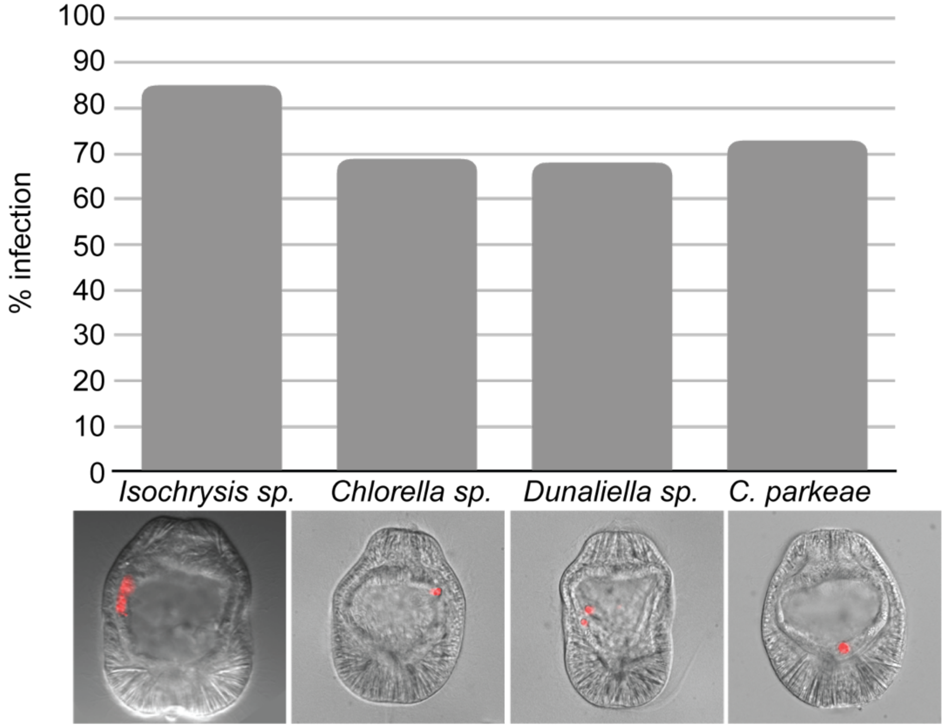


**Fig. S1**

Additional microalgae screened: *Isochrysis sp., Chlorella sp., D. salina, and C. parkaea. Aiptasia* larvae were infected at 4-6 days post fertilisation (dpf) for 24 hours and were washed into fresh FASW. Images are DIC and red autofluorescence of algal photosynthetic pigments.

### Fig. S2

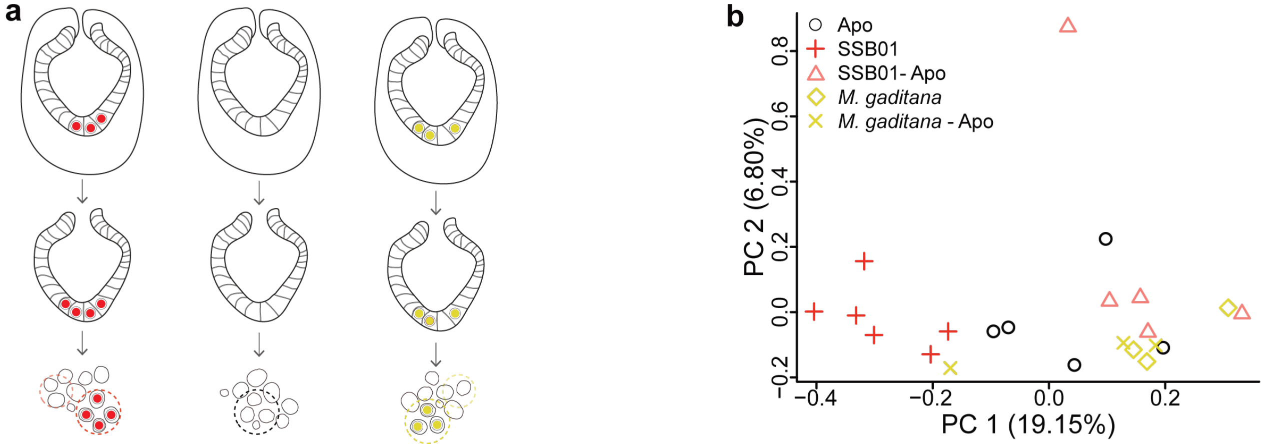


### Fig. S2

**a** Schematic of *Aiptasia* larvae used for cell-type specific sequencing. Ectodermal cells were removed resulting in only endodermal cells, which were dissociated and selected for based on contents: aposymbiotic cells from symbiotic larvae (SSB01-Apo), symbiotic cells from symbiotic larvae (SSB01), aposymbiotic cells from aposymbiotic larvae (Apo), cells containing *M. gaditana* from larvae infected with *M. gaditana* (Mg), and aposymbiotic cells from larvae infected with *M. gaditana* (Mg-Apo). **b** Principal Component Analysis (PCA) plot of host gene expression in different conditions.

**Fig. S3**

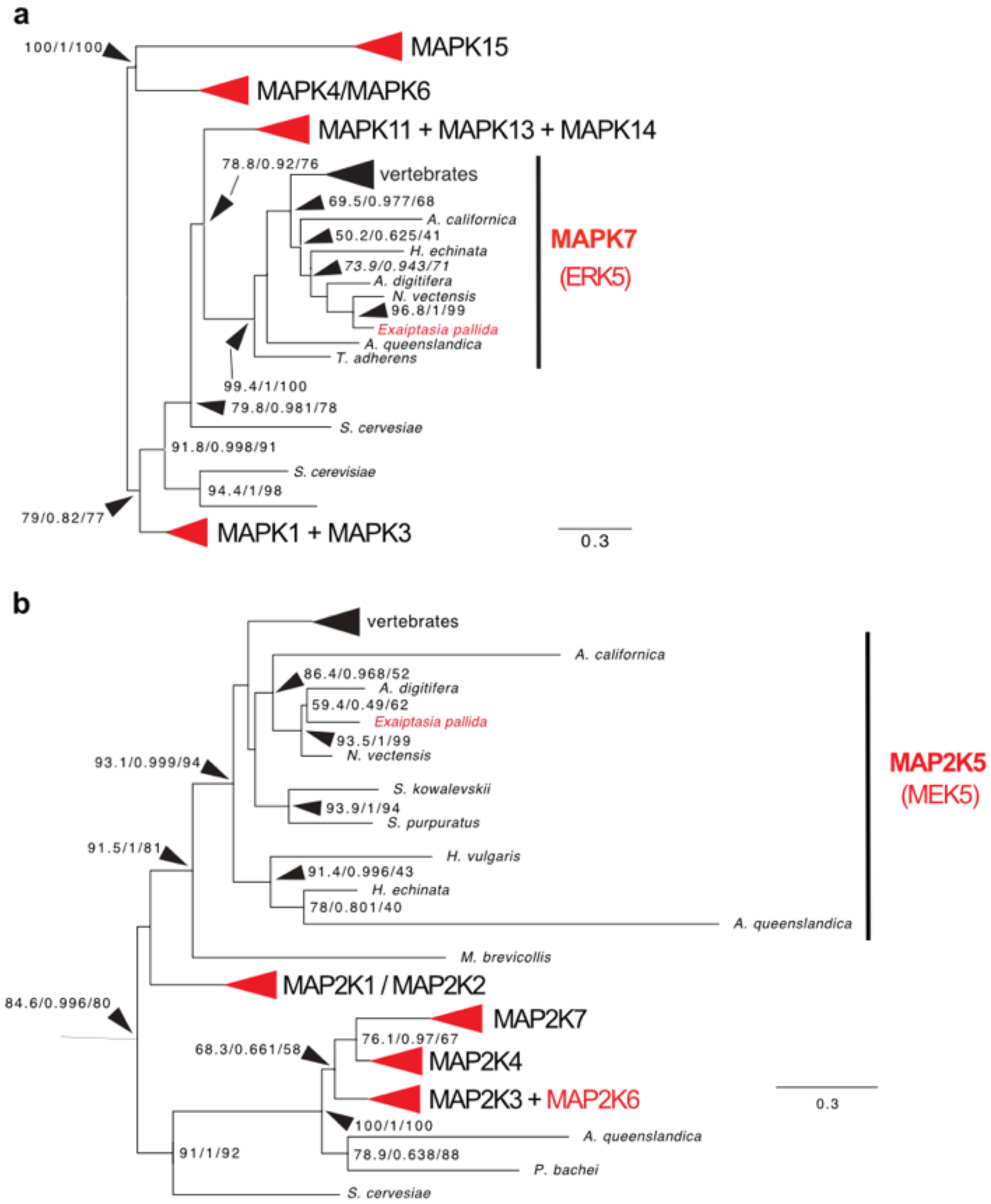


**Fig. S3**

Phylogenetic analysis of ERK5 and MEK5 from *Aiptasia*. **a** + **b** are collapsed trees of Aiptasia MAPK (a) or MAP2K (b) in comparison to several other cnidarian and vertebrate species. Red arrowheads or writing indicate presence of an *Aiptasia* homolog. Both *Aiptasia* ERK5 and MEK5 cluster within ERK5 (MAPK7) or MEK5 (MAP2K5), respectively. Full tree can be accessed through file S1 and S2.

### Table S1 – Statistics of transcriptional suppression of host cell immunity

Enumeration of modulation of innate immunity genes over 10 immune pathways, with some genes present in multiple pathways, and multiple transcripts annotated as the same gene (e.g. TRAF3)

| Pathway | Genes in Kegg-Pathway | Transcripts in Aiptaisa | SSB01 vs. Apo | SSB01 vs. SSB01-Apo | SSB01-Apo vs. Apo | *M. gaditana* vs. Apo | *M. gaditana* vs. *M. gaditana* -Apo | *M. gaditana* -Apo vs. Apo | SSB01 vs. *M. gaditana* | Unique count (KO-number) |
|---|---|---|---|---|---|---|---|---|---|---|
| C-type lectin receptor signaling pathway | 82 | 61 | 6 (9.84 %) | 8 (13.11 %) | 2 (3.28 %) | 2 (3.28 %) | 2 (3.28 %) | 2 (3.28 %) | 4 (6.56 %) | 29 |
| Complement and coagulation cascades | 78 | 14 | 0 (0 %) | 1 (7.14 %) | 0 (0 %) | 0 (0 %) | 0 (0 %) | 0 (0 %) | 0 (0 %) | 6 |
| JAK-STAT signaling pathway | 138 | 25 | 1 (4 %) | 4 (16 %) | 1 (4 %) | 1 (4 %) | 1 (4 %) | 0 (0 %) | 1 (4 %) | 16 |
| MAPK signaling pathway | 235 | 156 | 13 (8.33 %) | 25 (16.03 %) | 3 (1.92 %) | 1 (0.64 %) | 3 (1.92 %) | 2 (1.28 %) | 13 (8.33 %) | 92 |
| NF-kappa B signaling pathway | 89 | 45 | 10 (22.22 %) | 11 (24.44 %) | 3 (6.67 %) | 0 (0 %) | 3 (6.67 %) | 4 (8.89 %) | 6 (13.33 %) | 26 |
| NOD-like receptor signaling pathway | 139 | 117 | 14 (11.97 %) | 16 (13.68 %) | 1 (0.85 %) | 1 (0.85 %) | 5 (4.27 %) | 6 (5.13 %) | 6 (5.13 %) | 51 |
| RIG-I-like receptor signaling pathway | 53 | 41 | 5 (12.2 %) | 7 (17.07 %) | 1 (2.44 %) | 0 (0 %) | 3 (7.32 %) | 4 (9.76 %) | 2 (4.88 %) | 21 |
| TGF-beta signaling pathway | 79 | 44 | 3 (6.82 %) | 5 (11.36 %) | 3 (6.82 %) | 0 (0 %) | 1 (2.27 %) | 2 (4.55 %) | 3 (6.82 %) | 35 |
| TNF signaling pathway | 90 | 58 | 11 (18.97 %) | 10 (17.24 %) | 2 (3.45 %) | 4 (6.9 %) | 5 (8.62 %) | 7 (12.07 %) | 4 (6.9 %) | 33 |
| Toll-like receptor signaling pathway | 76 | 40 | 8 (20 %) | 8 (20 %) | 3 (7.5 %) | 1 (2.5 %) | 2 (5 %) | 4 (10 %) | 4 (10 %) | 23 |
| Unique transcripts within immune pathways | 744 | 386 | 36 (9.33 %) | 53 (13.73 %) | 11 (2.85%) | 6 (1.55 %) | 15 (3.89 %) | 15 (3.89 %) | 25 (6.48 %) | 207 |

### Table S2 – Live imaging statistics

| | # larvae imaged | total # of particles | mean # of particles | # expulsion events | # particle reacquisition | # larvae with replicating particles | % of larvae with expulsion event |
|---|---|---|---|---|---|---|---|
| SSB01 | 13 | 43 | 3.3 | 1 | 2 | 11 | 7.7 |
| *M. gaditana* | 7 | 19 | 2.7 | 5 | 5 | 0 | 57.1 |
| *N. oculata* | 7 | 13 | 1.9 | 12 | 11 | 0 | 87.5 |
| *C. velia* | 8 | 12 | 1.5 | 30 | 25 | 1 | 75 |
| Beads | 11 | 28 | 2.5 | 42 | 36 | 0 | 100 |

**Video S1**

3D reconstruction of LAMP1 staining in *Aiptasia* larvae infected with SSB01. LAMP1 is stained in magenta, DNA is stained with Hoechst in cyan, and autofluorescence of symbiont is shown in white.

**Video S2**
